# Diverging light-responsive behaviors in genetically similar strains of *Hydra vulgaris*

**DOI:** 10.64898/2026.11.18.767650

**Authors:** Gabriela Franco, Nahom Zerai, Matthew R. Gibson, Celina Juliano, Jacob T. Robinson

## Abstract

Sensory perception drives behavior by bridging the external environment with internal states. The freshwater cnidarian *Hydra vulgaris* serves as a useful model for capturing these dynamics due to its structural simplicity and distinct, state-dependent behaviors. However, few studies examine behavioral differences between strains that may arise from distinct genetic backgrounds, and none have compared behaviors from closely related lines. Here, we compare the phototactic profiles of two independently derived F1 lines created by self-crossing the standard laboratory AEP line: *Hydra vulgaris* AEP Irvine and AEP Kiel. While satiety alters behavior in both lines, starved AEP Irvine animals display stochastic wandering rather than the robust positive phototaxis seen in the AEP Kiel strain. A biased-correlated random walk model reproduces both behaviors by modifying jump rate and head angle. Despite lacking directional phototaxis, AEP Irvine remains light-sensitive: in darkness, animals exhibit fewer jumps, greater jump distance, and longer head-angle persistence. Behavioral motif classification reveals that light exposure alters time spent in stillness and elongation, as well as a few transition probabilities. These findings highlight behavioral differences between *Hydra* strains derived from the same parental line, providing an opportunity to identify molecular and genetic mechanisms coupling photoreception to state-dependent directional navigation.

## Introduction

Animal behavior is influenced by internal states that modulate responses to environmental cues. Organisms with simple, traceable nervous systems facilitate the discovery of fundamental relationships between molecular signaling, neural activity, and behavior. *Hydra vulgaris* possesses anatomical and neural simplicity, making it a powerful model organism for understanding state-dependent behaviors. *Hydra* consists of five major anatomical regions: the basal disk (i.e., foot), peduncle, body column, hypostome (i.e., head), and tentacles. The animal contains two tissue layers with separate nerve nets and 15 transcriptionally distinct neuronal subtypes (Little et al., 2025). These neurons organize into contraction burst, rhythmic potential 1, and rhythmic potential 2 circuits, which are responsible for distinct behavioral repertoires, such as contraction, elongation, swaying, and somersaulting locomotion (Dupre & Yuste, 2017; Han et al., 2018; Yamamoto & Yuste, 2023).

The diffuse nerve net of *Hydra* integrates environmental inputs through neural processing to produce motor responses. Previous work established that *Hydra* demonstrates sensorimotor reflexes to thermal (Bosch et al., 1988; <u>Mast, 1903; Schroeder & Callaghan, 1981</u>), chemical (Badhiwala et al., 2018; Kepner & Hopkins, 1924; Lenhoff, 1961; Scappaticci et al., 2010), mechanical (Badhiwala et al., 2021; Mast, 1903; Rushforth et al., 1963; Wagner, 1905), and optical stimuli (Giez et al., 2024; Kim et al., 2024; Passano & McCullough, 1962; Plachetzki et al., 2012; Rushforth et al., 1963; Singer et al., 1963; Trembley, 1744). Specifically, the oral region (containing hypostome and tentacles) is known to process sensory information (Badhiwala et al., 2021).

An example of sensorimotor integration is positive phototaxis, in which *Hydra* utilizes directional somersaulting to navigate toward a light source. Recent work quantified this behavior in *Hydra vulgaris*, demonstrating that satiety modulates positive phototaxis (Kim et al., 2024). Furthermore, satiety regulates gene expression, including opsins and neuropeptides, within specific neural circuits, such as Ec3 (N3) and En1 (N4) networks (Giez et al., 2024). However, a major limitation of these behavioral studies is the assumption that all behaviors are uniform across *Hydra* strains, overlooking how phenotypes can vary even between closely related genetic animals.

The studies described here began with the observation that closely related strains of *Hydra vulgaris* cultured under identical conditions show different phototactic behavior following starvation (>10 days). Specifically, the *Hydra vulgaris* AEP Kiel strain migrated toward the incubator’s light source (Kim et al., 2024), whereas the *Hydra vulgaris* AEP Irvine strain did not. These two lines were both derived from self-crossing the parental *Hydra vulgaris* AEP strain (PMID: 16556723) (see materials and methods for details). Given the close genetic relationship between these strains, we were intrigued by their divergent behavior and set out to identify which specific aspects of their behavior varied and how those differences could alter phototaxis. Building on the phototaxis assay described in Kim et al. (2024), we quantified somersaulting (“jump”) frequency, jump distance, and head orientation across both strains. We demonstrated that differences in jump rate and head alignment modeled these trajectories within a Biased Correlated Random Walk (BCRW) framework. Finally, to determine how light affects the AEP Irvine strain, we compared behavior across light and dark conditions. We found that light modulates motor output, such as jump rate and jump distance. Furthermore, using a behavioral state classifier, we found that while time spent in contraction was consistent between light and dark conditions, somersaults, stillness, and elongation occupancy differed, as did the likelihood of transitioning from contraction to either elongation or stillness. To our knowledge, this study is the first to report behavioral differences between strains of *Hydra vulgaris* under the same environmental cue and internal states. Combining these behavioral models with comparative genomics and transcriptomics, and neural circuit mapping will provide a foundation for predicting how subtle molecular and circuit-level variations dictate behavioral states.

## Results

### Divergent Phototactic Behavior in Genetically Similar F1 *Hydra* Lines

We compared the phototactic profiles of two independently derived lines sharing an AEP lineage by using the high-throughput imaging platform under a directional LED optical gradient (**Figure 1A**). To directly compare phototactic behaviors between *Hydra* strains, we reanalyzed datasets from the AEP Kiel strain (**Figure 1B–D**; Kim et al., 2024) and evaluated them together with our newly collected tracking data from the AEP Irvine strain (**Figure 1E-G**). As Kim et al. reported that behaviors depend on satiety state, we tested both fed (fed 2 days prior to experiment) and starved (fed 2 weeks prior to experiment) conditions. We focused the analysis on the initial 8 h window, or until the foot position was within 7 mm of the light source to complete positive phototaxis.

**Figure 1.**
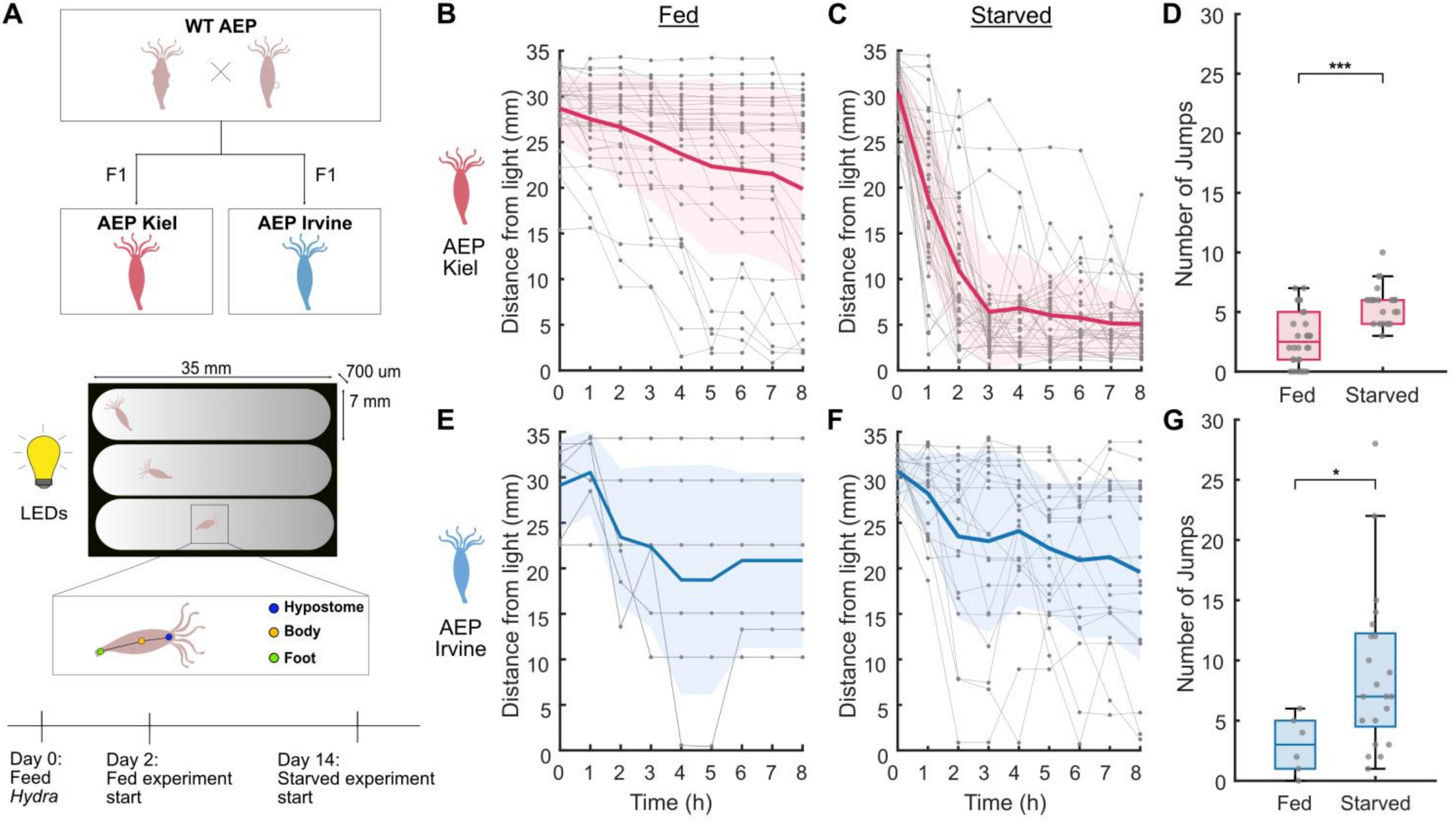
Distinct phototactic behavior in *Hydra* lines derived from the AEP background. **(A)** *Top:* Schematic illustrating the relationship between *Hydra vulgaris* strains: AEP Irvine and AEP Kiel are independently derived F1 lines originating from a parental AEP strain. *Bottom:* Experimental set-up and timeline. *Hydras* were inserted into a 35 mm phototaxis chamber with directional LED illumination on one side and recorded at 1 frame per second (fps). **(B, C)** Foot position of animals at every hour for 8 h (t = 0, 1, …, 8 h) relative to the light boundary (mm) for **(B)** fed AEP Kiel (2 days post-feeding; n=29), and **(C)** starved AEP Kiel (2 weeks post-feeding; n=24; data adapted from Kim et al., 2024). Bold line and shaded band indicate mean ± SD, and in gray are individual traces. **(D)** Total jumps in the first 8 h or until positive phototaxis in the fed (n=29) and starved AEP Kiel (n=24). Fed AEP Kiel displayed fewer jumps than starved (p < 0.001; Mann-Whitney U Test). **(E, F)** Foot position of animals at every hour for 8 h (t = 0, 1, …, 8 h) from the light boundary (mm) for **(E)** fed AEP Irvine (2 days post-feeding; n=6) and **(F)** starved AEP Irvine (2 weeks post-feeding; n=21). **(G)** Total jumps in the first 8 h or until positive phototaxis in the fed (n=6) and starved AEP Irvine (n=21). Fed AEP Irvine displayed fewer jumps than starved (p = 0.015; Mann-Whitney U Test). All box plots show medians, first, and third quartiles.

Comparing positions between AEP Kiel and AEP Irvine animals across fed and starved conditions revealed that strain-specific differences emerge when starved. In the fed condition, neither strain had a directional preference toward the light source at 8 h, as both AEP Kiel (19.9 ± 10.3 mm) and AEP Irvine (20.9 ± 9.6 mm) exhibited dispersed positions that averaged near the center of the chamber (**Figure 1B, E**). However, in the starved condition, the AEP Kiel animals demonstrated a robust migration toward the light source, while starved AEP Irvine did not display the same directed movement (**Figure 1C, F**). At 8 h, starved AEP Kiel animals were near the light source (5.0 ± 4.0 mm), while the starved AEP Irvine animals were dispersed throughout the imaging chamber (16.6 ± 9.4 mm). This observation persisted until the end of the recording at 48 h (**Figure 1 – figure supplement 1**).

Despite similar positions at 8h between fed and starved AEP Irvine animals, we found that satiety modulates the number of jumps in both strains. Starvation induced a 2.4-fold increase in total jumps in AEP Kiel (p < 0.001; Mann-Whitney U Test), and a 2.3-fold increase in AEP Irvine (p = 0.015). (**Figure 1D, G**). This demonstrates that while food deprivation increases jumps in both strains, it only results in directed phototaxis in the AEP Kiel strain, and undirected exploration in the AEP Irvine strain.

Having established that starvation induces somersaulting, we next evaluated kinematic parameters to identify additional strain-dependent differences in starved strains. We found that the behavioral divergence between strains stems from jump frequency and directional coordination, rather than underlying somersaulting mechanics. For instance, jump distance was highly consistent between strains, indicating that both lines execute comparable somersaults (p = 0.14, Mann–Whitney U test; **Figure 2A**). However, the two strains exhibited a difference in jump frequency, as the AEP Kiel strain displayed a significantly higher jump rate compared to AEP Irvine (p < 0.001, Mann–Whitney U test; **Figure 2B**), establishing AEP Kiel as the more active line upon light exposure. Furthermore, jump direction analysis revealed that Kiel executed significantly more jumps toward the light source than away from it (p < 0.001, Mann–Whitney U test; **Figure 2C**, left). Although AEP Irvine also showed a preference for jumping toward light (p =0.011; **Figure 2C**, right), their trajectory was far more variable, resulting in reduced net migration.

**Figure 2.**
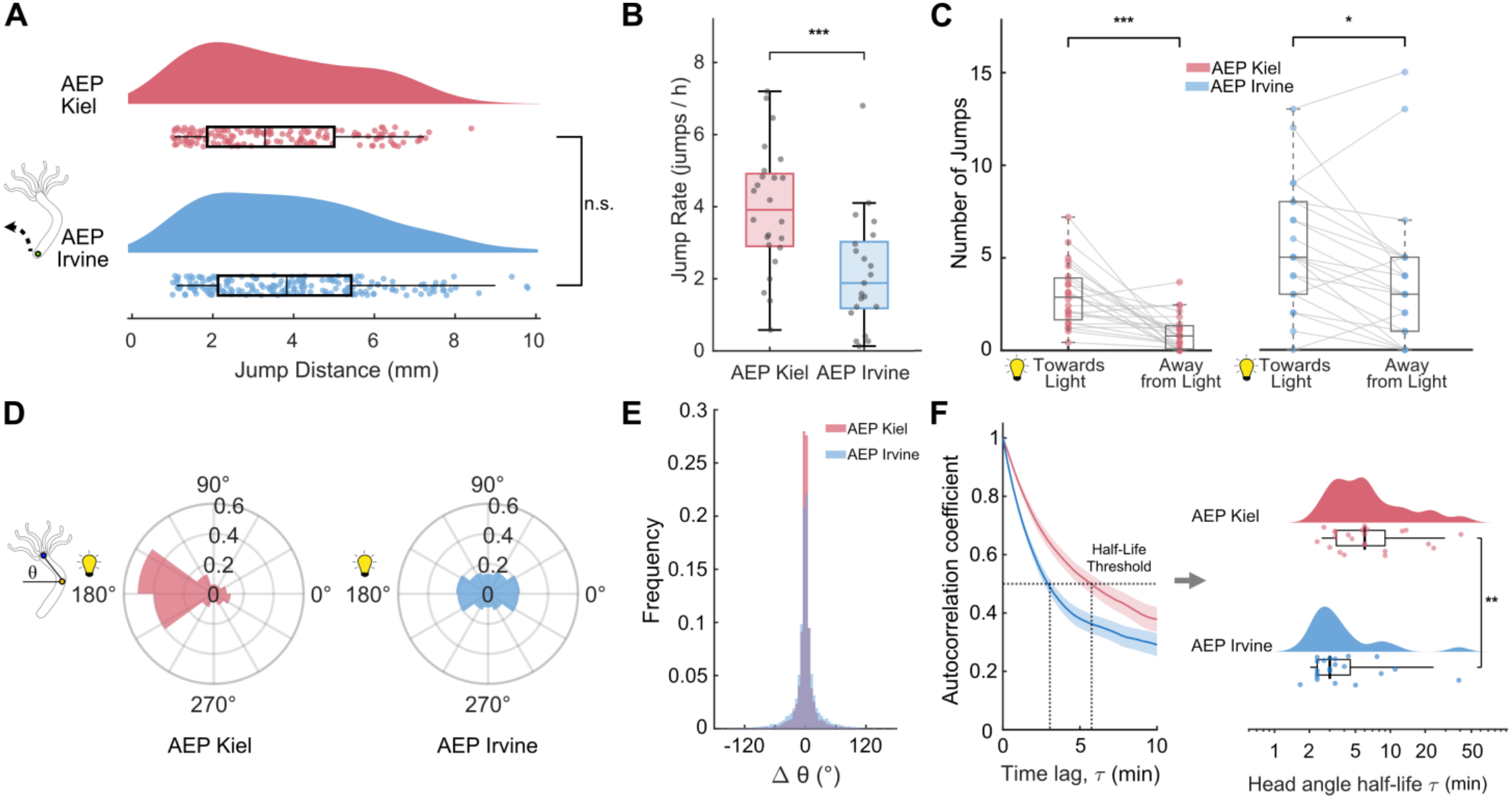
Strain-dependent differences in kinematic profiles between *Hydra vulgaris* AEP Kiel and AEP Irvine. **(A)** Distribution of jump distances (mm) in AEP Kiel (red, n = 24) and AEP Irvine (blue, n = 21) strains. AEP Kiel jump distances and AEP Irvine are comparable (p = 0.14, Mann–Whitney U test). **(B)** Jump rates between strains. AEP Kiel animals exhibit higher jump rate than AEP Irvine (p < 0.001, Mann–Whitney U test). **(C)** Directional preference of individual jumps oriented “Towards Light” versus “Away from Light” for Kiel (left) and Irvine (right). Kiel displayed strong preference towards light vs. away from light (p < 0.001, Wilcoxon signed-rank test). AEP Irvine also displayed directional bias toward light vs. away from light (p = 0.011, Wilcoxon signed-rank test). **(D)** Angular head orientations relative to the light source (180°) with AEP Kiel (red) and AEP Irvine (blue). **(E)** Distribution of changes in head angle (ΔΘ). **(F) Left:** Head angle autocorrelation coefficient C(τ) versus time lag τ; horizontal line denotes the half-life threshold (0.5). Bold line and shaded band indicate mean ± SEM. **Right:** Log-scale comparison of behavioral half-lives plotted as median and interquartile-range (p = 0.003, Mann–Whitney U test). All box plots show medians, first, and third quartiles.

This pattern was further reflected in the head angle orientation and persistence of the two strains. The AEP Kiel strain maintained a unidirectional head angle (*θ*), with the head focused on the light source (**Figure 2D**, left). Conversely, the AEP Irvine strain exhibited a broad head angle distribution, indicating the absence of a target orientation (**Figure 2D**, right). However, changes in head angle (Δ*θ*) were centered around 0° for both strains (**Figure 2E**). The two strains diverged again in how long they held a given head angle. To evaluate head angle persistence, we calculated the half-life of the circular autocorrelation decay, which quantifies the stability and periodicity of head-sweeping over time (**Figure 2F**). AEP Kiel demonstrated prolonged persistence, maintaining its orientation with a longer half-life (τ_1/2_) of 4.3 min (IQR = 3.5–5.3), whereas for AEP Irvine, the head angle decorrelated faster, dropping τ_1/2_ to 3.0 min (IQR = 2.3–4.1; p = 0.003, Mann-Whitney U-Test; **Figure 2F**). This loss of correlation confirms that AEP Irvine exhibits frequent head swaying rather than maintaining a fixed angle. While AEP Irvine does not show strong directional phototaxis like AEP Kiel, it actively responds to light through exploration and frequent head adjustments.

### Computational Modeling of Divergent Phototactic Behaviors Using Biased Correlated Random Walks

Having identified clear strain-dependent differences in jump rate and head orientation, we next tested whether these kinematics were sufficient to mathematically explain the behavioral phenotypes of the AEP Kiel and Irvine strains. To do so, we constructed a Biased Correlated Random Walk (BCRW) using three derived parameters unique to each strain: turning angle (Δ*θ*), hourly jump frequency (*λ*), and jump distance (*d*). By incorporating a relative weighting parameter (*w*) that balances persistence with directional bias, we simulated trajectories over an 8 h window (**Figure 3 – figure supplement 1**). We then evaluated whether these simulations could replicate the behavioral divergence observed between strains (**Figure 3**).

**Figure 3.**
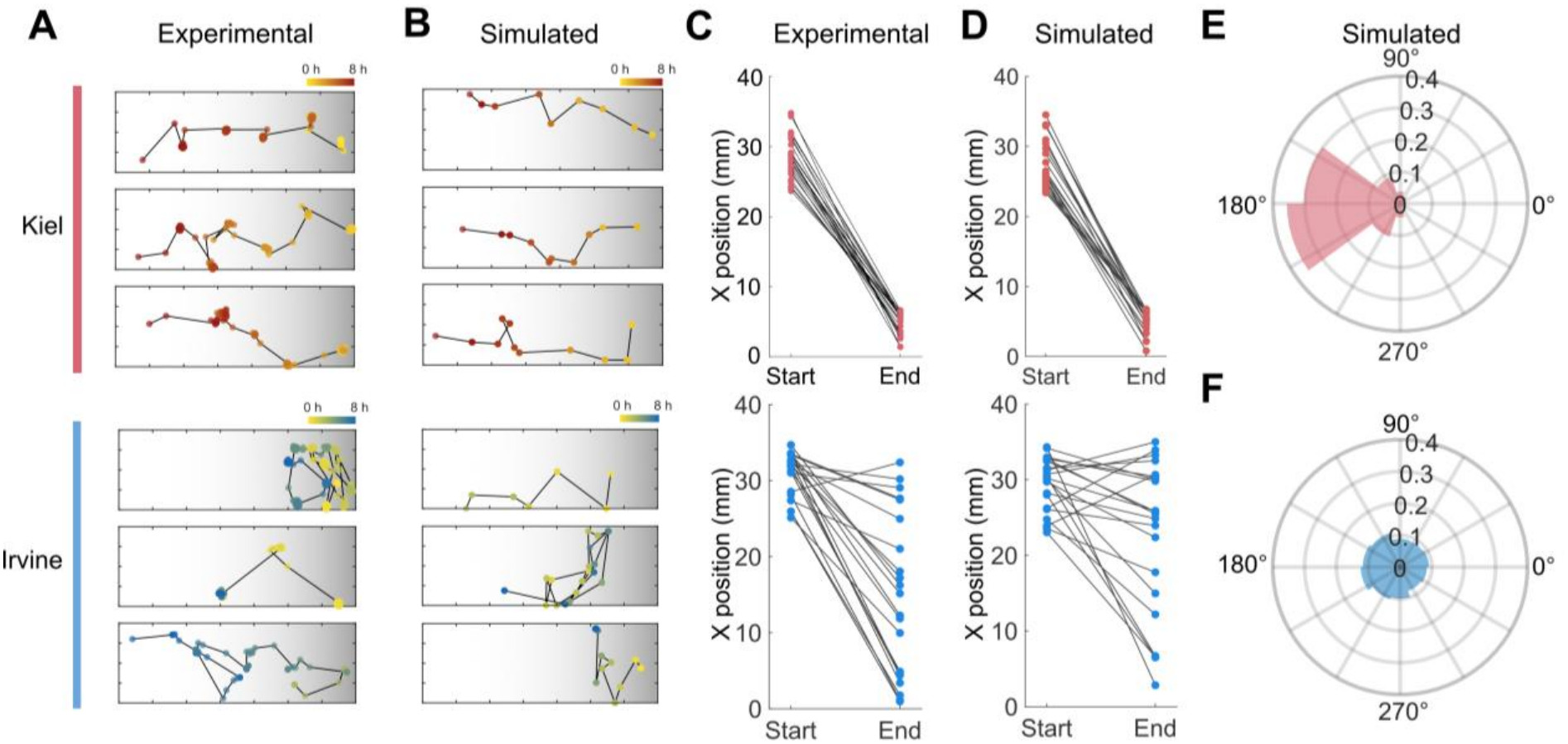
Computational modeling replicates BCRW dynamics of *Hydra vulgaris* AEP Kiel and AEP Irvine strains using head angle, jump rate, and jump distance parameters. **(A, B)** Representative foot traces comparing **(A)** experimental tracking data with **(B)** simulated paths derived from model parameters for the AEP Kiel (n = 24; top panels, red/orange lines) and AEP Irvine (n = 21; bottom panels, blue/green lines) strains. Marker colors indicate time progression from start to 8 h against a horizontal light gradient background (darker gradient denotes proximity to the light source). **(C, D)** Population-level displacement trends over the 8 h window, tracking the horizontal X-axis position (mm) from start to end for **(C)** experimental animal groups and **(D)** corresponding simulation. **(E, F)** Polar plots showing the angular head orientation distributions generated by the **(E)** simulated AEP Kiel model and **(F)** simulated AEP Irvine model under directional illumination (light positioned at 180°).

We found that by adjusting only the jump rate and head angle variables in our model, we could accurately simulate the different trajectories observed in the AEP Kiel and AEP Irvine locomotion (**Figure 3A, B**). In the AEP Kiel strain, both experimental observations and computational simulations reanalyzed and simulated from Kim, et al. (2024) depicted direct, progressive pathways toward the light (**Figure 3A, B**, top panels). Conversely, experimental and simulated trajectories for the AEP Irvine strain produced high spatial scattering and lack of directional migration, reflecting a highly variable and uncoordinated spatial search pattern (**Figure 3A, B**, bottom panels). Furthermore, the BCRW modeled population-level displacement trends (**Figure 3C, D**). The simulated AEP Kiel closely mirrored the experimental animals, demonstrating a highly coordinated directional migration from their initial starting positions to their final destinations near the light source. Similarly, the simulated AEP Irvine accurately produced end positions scattered across the chamber as observed experimentally.

The model successfully captured the head angle dynamics unique to each lineage under directional illumination (**Figure 3E, F**). The simulated AEP Kiel generated an angular head distribution oriented towards the 180° light source (**Figure 3E)**, driven by a high directional bias weight (**Table 1**). Meanwhile, the simulated AEP Irvine produced a broad, omnidirectional distribution without a preferred target angle (**Figure 3F; Table 1**). Minor stochastic variations between the experimental and simulated profiles are expected due to the probabilistic nature of the random walk algorithm. Overall, these simulations demonstrate that head angle directionality and jump rates simulate distinct, strain-specific phototactic phenotypes in a BCRW framework.

**Table 1.** Empirical and simulated parameters for the Biased Correlated Random Walk (BCRW) model.

| **Strain** | **Experimental Bias (R<sub>bias</sub>)** | **Experimental Persistence (R<sub>turn</sub>)** | **Experimental Weight (w)** | **Simulated Bias (R<sub>bias</sub>)** | **Simulated Persistence (R<sub>turn</sub>)** | **Simulated Weight (w)** |
|---|---|---|---|---|---|---|
| **AEP Kiel** | 0.4644 | 0.9491 | 0.3286 | 0.6597 | 0.9368 | 0.4132 |
| **AEP Irvine** | 0.0888 | 0.9130 | 0.0886 | 0.0991 | 0.9174 | 0.0975 |

### *Hydra vulgaris* AEP Irvine is light–responsive and has altered behavioral responses

Having validated that BCRW simulations capture phototactic trajectories, we next asked whether light actively modulates AEP Irvine behavior or if its exploration is completely light independent. To determine whether light acts as a sensory input in AEP Irvine, we compared spatial distributions and characterized its movement under light versus dark conditions.

Starting AEP Irvine animals on the side closest to the light-source produced foot distributions like those started on the far side, indicating that initial orientation relative to the light source does not change the final foot position (16.6 ± 9.4 mm, **Figure 1F**; 12.4 ± 7.1 mm, **Figure 4A, left**). Similarly, in the dark, AEP Irvine animals exhibited a dispersed spatial distribution over time, with the final positions near the chamber center with high individual variability (10.1 ± 8.4 mm, **Figure 4A, right**). A two-way mixed ANOVA confirmed that while overall spatial distributions shifted over time, there was no effect of the light condition or light-time interaction. Together, these results demonstrate that AEP Irvine spatial positioning is unaffected by directional light, while also ruling out potential responses to thermal gradients generated by the light source (**Figure 4–figure supplement 1**).

**Figure 4.**
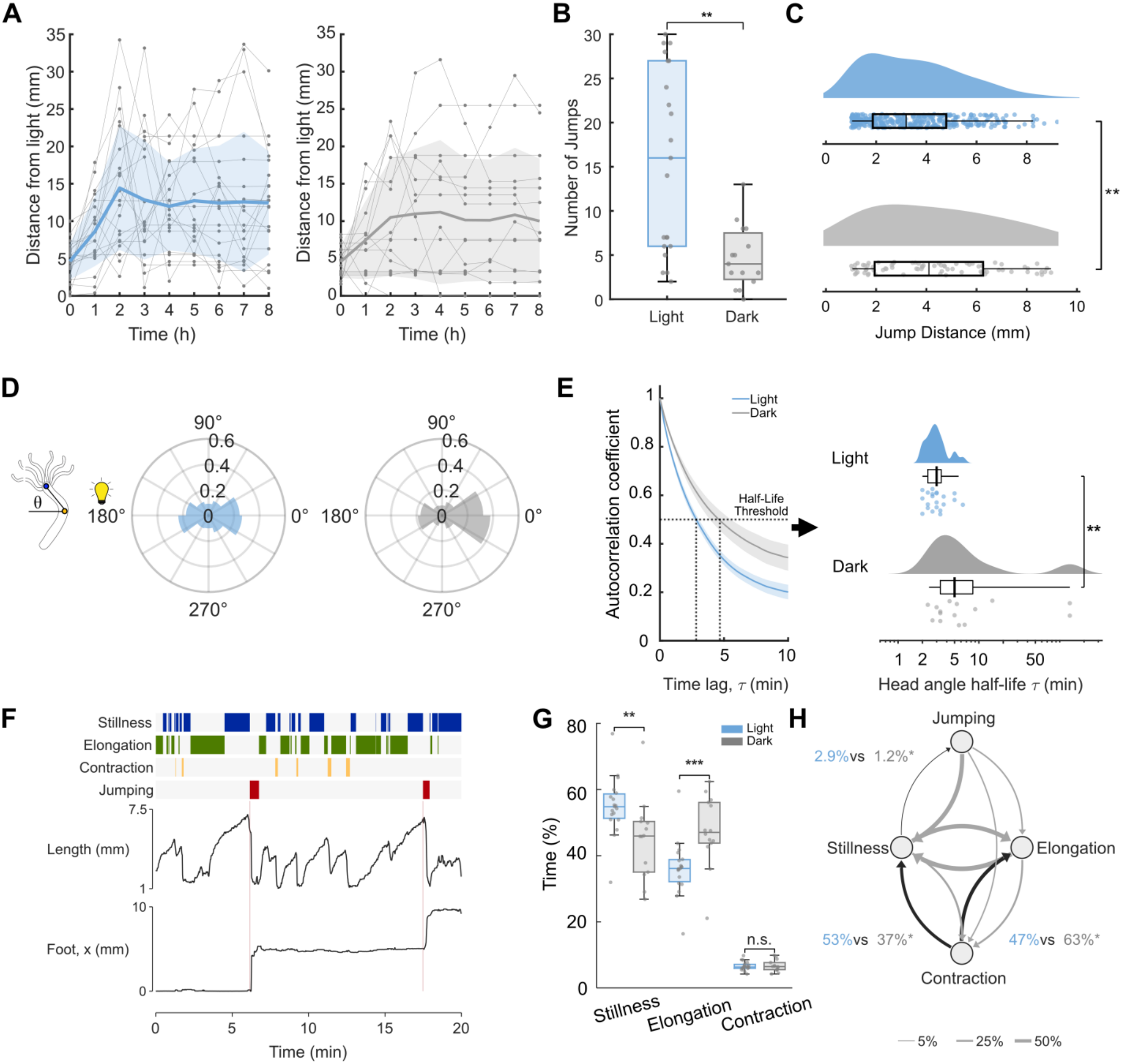
*Hydra vulgaris* AEP Irvine strain exhibits light-dependent responses in behavior. **(A)** Foot locations sampled every hour across an 8 h timeline (t = 0, 1, …, 8 h) relative to the light boundary under directional light (blue curve, left, n = 21) and blocked light (grey curve, right, n = 15) conditions. Bold line and shaded band indicate mean ± SD, and in gray are individual traces. Spatial positions shiftzed over time (F3.3,114 = 7.19; p < 0.001), but positions did not differ between conditions (Main effect of light: F_1,34_ = 1.08; p = 0.30; Light x Hour: F_3.3,114_ = 0.42; p = 0.76). **(B)** Total number of jumps under light versus control conditions (p = 0.0014, Mann–Whitney U test). Light significantly increases overall jump rates. **(C)** Jump distance distributions for starved animals under light (top) and control (bottom) conditions (p = 0.002, Mann–Whitney U test). **(D)** Angular head orientation (θ) relative to the light source (positioned at 180°). Light-exposed animals (left) and control animals (right). **(E)** Left: Head angle autocorrelation coefficient versus time lag; the horizontal dashed line denotes the half-life threshold (0.5). Right: Log-scale comparison of calculated head angle half-life (τ_1/2_), showing half-life (p < 0.001, Mann–Whitney U test). **(F)** Representative sample of classification of behavioral events. **Top:** Time-series of classified behaviors. **Middle:** Example kinematic input feature (body length). **Bottom:** Foot position along the x-axis, with jumps labeled in pink. **(G)** Distribution of time spent (%) on behaviors for light versus dark control conditions, where light animals are more still than dark control (p = 0.002, Mann–Whitney U test), light animals are less elongated than dark control (p < 0.001). Contraction is similar (p = 0.96). **(H)** Behavioral state transition diagram. Light animals go to stillness more often: 53% vs 37% (p < 0.001, Mann–Whitney U test). Light animals go to elongation less often 47% vs 63% (p < 0.001). Jump follows more than twice as often in light, 2.9% vs 1.2% (p = 0.013). All box plots show medians, first, and third quartiles.

When comparing AEP Irvine jump behaviors between light and dark conditions, we found that the number of jumps and jump distance changes in the presence of light. Despite similar spatial distributions, the number of jumps increased 4-fold in the light compared to the dark control. This indicates that light stimulates the *Hydra* and results in more somersaulting (**Figure 4B**). Interestingly, the increased jump frequency is inversely correlated with the size of the jump, as the distribution of jump distances revealed smaller jump sizes under directional light exposure (3.0 mm, IQR = 1.76–4.62) compared to the larger distances observed under dark control conditions (4.1 mm, IQR = 1.94–6.26; **Figure 4C**). Together, these metrics demonstrate that while AEP Irvine lacks directional phototaxis, light exposure induces a highly active yet spatially undirected behavioral phenotype, characterized by frequent, short-range displacements.

In addition to jump kinematics, we unexpectedly observed differences in head angles between the light and dark conditions, as animals in the dark had a bias due to proximity to the walls rather than directional steering (**Figure 4D**). While light-exposed animals showed no apparent head angle preference, pointing randomly across all angles (*θ*) relative to the light source (positioned at 180°), dark animals had majority distribution toward the opposite end of the chamber at 0°, (**Figure 4D**). Because head orientation aligns with the direction of the jump, we further analyzed animals in the dark if they displayed a low number of jumps (< 5 jumps) and high number of jumps (≥ 5 jumps) (**Figure 4–figure supplement 2**). AEP Irvine *Hydra* with a high number of jumps exhibited a uniform head angle distribution with no preferred direction, comparable head angle orientations to the light condition animals. Animals with low amounts of jumps displayed a bias pointing away from their initial placement site. The physical boundary constraints near the chamber wall combined with the lack of locomotion reflect localized positioning rather than active directional steering.

In addition, light distinctly reduced head angle persistence. To quantify this, we measured the decay half-life of circular autocorrelation (τ_1/2_; **Figure 4E, left**). Light exposure accelerated the decorrelation rate, reducing the median half-life from 5.0 min (IQR = 3.33–8.42) in dark controls to 3.0 min (IQR = 2.33–3.42) under light (p < 0.001, Mann–Whitney U test; **Figure 4E, right**). This loss of correlation confirms that light stimulates head-sweeping. Therefore, while AEP Irvine does not show directional phototaxis, it actively responds to light through increased exploration and frequent head adjustments.

To understand how light influences behavioral repertoire, we constructed a classifier to identify and quantify discrete behaviors and evaluate their transition probabilities. Specifically, we identified three distinct behaviors using an L1-regularized logistic regression model (stillness, elongation, and contraction), while a fourth behavior, jumping, was labeled manually. We built on prior work that used machine learning to identify *Hydra* basal behaviors and modeled our states after the six behaviors documented by Han et al. (2018). However, we omitted tentacle sway due to video resolution limitations, along with bend/sway as its classification performance was slightly above the shuffled null baselines (AUC 0.74 vs 0.66; **Figure 4–figure supplement 3**).

Pose estimation coordinates for the head, body, and foot were extracted at 1 s resolution and filtered for low-confidence detections prior to behavioral state classification (**Materials and methods**). We trained the classifier using eight kinematic features derived from tracking coordinates. Model performance transferred effectively from our training to our testing animals starting towards the light source (AUC 0.77– 0.97; **Figure 4–figure supplement 3**), resulting in continuous sequences of behaviors (**Figure 4F**).

Across light conditions, AEP Irvine exhibited significant differences in how they divide their time across the distinct repertoire **(Figure 4G)**. Animals exposed to light spent more time in stillness than those in the dark (median 55% vs 46%; p = 0.002), and less time elongating (36% vs 47%; p < 0.001), while time spent contracting did not differ between groups (6.3% vs 6.5%, p = 0.96). Light exposure also changed transitions between certain behaviors (**Figure 4H**). Following contractions, *Hydra* in the light transitioned towards stillness more frequently (53% vs 37%; p < 0.001) and correspondingly transitioned less toward elongation (47% vs 63%; p < 0.001). Animals under light exposure also transitioned from stillness to jumping more than twice as often (2.9% vs 1.2% of stillness exits, p = 0.013). All other transitions were similar between conditions (p ≥ 0.18). Altogether, this analysis reveals that light stimulation reconfigures *Hydra* behavioral repertoire from consistent elongation to increased stillness and jumping.

## Discussion

Our observation that phototaxis in two genetically similar *Hydra vulgaris* strains under identical conditions display different phototactic behaviors raises the question of how subtle genetic variations alter sensory-motor processing within a simple nervous system.

A central insight from our kinematic analysis is the separation between biomechanical execution and directional steering. Both AEP Kiel and AEP Irvine exhibit identical jump step sizes (**Figure 2A**) and matching frequency distributions of angular head sweeping (**Figure 2E**), but the head orientation of AEP Kiel is directed towards the light source while AEP Irvine is not. These findings demonstrate that the biomechanics of jumping and head turning remain conserved, indicating that behavioral divergence stems from directional steering rather than motor deficits. The AEP Kiel strain exhibits phototaxis using light cues for directional steering, while the AEP Irvine strain utilizes stochastic wandering. The alignment between experimental and simulated pathways confirms that *Hydra* does not require high-level path integration or long-term spatial memory. Instead, phototaxis emerges as a direct mathematical consequence of persistent directional bias superimposed on localized somersaulting kinetics.

The origin of this altered directional bias points directly to underlying differences in the sensory-motor processes of *Hydra*. Lacking specialized visual organs, *Hydra* relies on a family of opsins distributed throughout its ectoderm and nervous system for photoreception (Macias-Muñoz et al., 2019; Plachetzki et al., 2012). In *Hydra*, starvation is linked to changes in gene expression, including opsins and neuropeptides (Giez et al., 2024), but the precise mechanisms on how starvation induces changes in phototaxis on a molecular level are unknown. Genomic divergence and strain-specific transcriptomic profiles across *Hydra* lines (Cazet et al., 2023; Chapman et al., 2010) may alter baseline opsin expression levels (Macias-Muñoz et al., 2019), thereby tuning light perception prior to downstream neural processing. Although the genetic basis of the behavioral differences is unknown, future studies using comparative transcriptomics and genetic mapping across AEP derivative lines could identify the responsible molecular pathways.

Although AEP Irvine foot locations appear similar in light versus dark conditions **(Figure 4A)**, we find key behavioral differences that occur across these two conditions, indicating that the light stimulus does modulate a subset of behaviors (**Figure 4B–H)**. AEP Irvine retains biomechanical execution, but exhibits altered jump distances, light-activated somersaulting, and reduced head persistence. Furthermore, AEP Irvine exhibits more quiescence in the light and appears as though to rest after contractions or before preparing to jump (**Figure 4G–H**). The AEP Irvine contrasts this in the dark, which executes significantly fewer somersaults, but has elevated jump sizes and elongations. Importantly, this demonstrates that the behavioral variation observed in AEP Irvine is driven by a divergence in how external light cues interact to guide spatial orientation when starved. While this strain does not consistently phototax in either fed or starved condition, these subtle differences in behavior provide insights into how light modulates behavior in *Hydra*, suggesting that neural circuits switch between active exploration and less active behavior.

From an ecological perspective, phototactic variability within *Hydra* populations may serve as a vital survival mechanism. In clonal and genetically similar organisms, behavioral divergence can aid populations against environmental fluctuations (Honegger & De Bivort, 2018; Moiron et al., 2020). Such individuality frequently emerges despite near-identical genetic backgrounds and rearing environments, as demonstrated in the clonal fish *Poecilia formosa* (Bierbach et al., 2017) and across developmental stages in *Caenorhabditis elegans* (Harel et al., 2024). In *Drosophila melanogaster*, this behavioral individuality is driven by stochastic variations during neural circuit assembly (Buchanan et al., 2015; Linneweber et al., 2020), with the degree of non-genetic variance itself operating under distinct genetic control (Ayroles et al., 2015). Although non-genetic behavioral diversity remains understudied in *Hydra*, wild *Hydra* are known to use bet-hedging strategies in reproductive contexts; for instance, embryos exhibit highly variable, sporadic hatching times to ensure population persistence in freshwater habitats (Steele et al., 2019). This suggests that individuality is plausible in *Hydra* as well, but more research is needed.

In summary, our findings establish that laboratory lines of *Hydra vulgaris* can evolve distinct behavioral light preferences while retaining identical locomotor mechanics. This finding that closely related *Hydra* strains have distinct behaviors, combined with a quantitative framework for tracking and analyzing behavior, offers an opportunity to investigate the neural and genetic mechanisms governing sensorimotor processing in *Hydra* and how subtle molecular and genetic differences can affect cnidarian behavior.

## Materials and Methods

### *Hydra vulgaris* strain and maintenance

Both lines used in this study descend from the parental *Hydra vulgaris* AEP strain (PMID: 16556723). Like all *Hydra* strains, AEP is maintained primarily through asexual budding but retains the capacity for sexual reproduction: individual polyps differentiate as either male or female and crossing them produces embryos that develop into F1 animals. The two F1 lines used here were each produced by self-crossing of the original AEP strain in independent laboratories: one in the Bosch laboratory (Kiel, Germany), hereafter AEP Kiel, and one in the Steele laboratory (Irvine, CA), hereafter AEP Irvine. Each line was founded from a single hatched F1 polyp and subsequently maintained by asexual budding. The Irvine line has also been referred to as SS1, an informal designation reflecting its elevated rate of gametogenesis. These lines differ in their phototactic behavior: AEP Kiel migrated toward the incubator’s light source, as we previously reported (Kim et al., 2024), whereas AEP Irvine did not.

The animals were cultured according to a protocol adapted from the laboratory of Robert Steele (University of California, Irvine, USA). Hydras were maintained at 18°C in an incubator on a 12 h:12 h light:dark cycle and were fed 3 times a week with freshly hatched *Artemia* nauplii (Brine Shrimp Direct). *Hydra* were not fed for 2 days prior to the experiment in the fed group to allow complete digestion, and for 14 days in the starved group. Animals that had gametes, buds, or appeared unhealthy (short tentacles, small size) were excluded from the analysis.

### Microfluidic device fabrication and usage

Microfluidic device fabrication and usage were performed as previously described by Kim et al. (2024). The microfluidic devices were fabricated using 9:1 polydimethylsiloxane (PDMS; Sylgard 184) with a resin 3D-print of 35 mm x 7 mm x 700 μm chambers serving as the mold. After curing the PDMS, ports on either side of each chamber were punched with a 1.25 mm diameter biopsy punch. The PDMS was O_2_ plasma-bonded to a 75 mm × 50 mm glass slide (Corning 2947-75X50). A single *Hydra* polyp was loaded into each chamber of the device using a syringe and tubing following a previously established protocol through the ports (Badhiwala et al., 2018; Tzouanas et al., 2021). The surface of the device was covered with optically clear adhesive well plate seals (Thermo Fisher Scientific AB1170) to minimize evaporation through the ports. *Hydra* were removed after each experiment. The devices were then cleaned by flushing with deionized (DI) water, 70% ethanol, and then DI water. The devices were soaked in DI water and boiled at 150°C for ∼2 h, and completely dried on hot plates until future use.

### Imaging setup and phototaxis assay

Behavioral tracking under directional light was performed using the microfluidic phototaxis platform and imaging protocol established by Badhiwala et al. (2018) and Kim et al. (2024). Briefly, individual animals were placed into customized chambers positioned on an acrylic platform. Directional illumination was supplied by an LED panel backed by an optical diffuser, generating a uniform optical gradient across the chamber with a peak power of ∼4.15 mW at the illuminated edge. For dark control experiments, uniform dim backlighting (R:G:B 1:1:1) was used to enable video tracking without providing a directional cue.

Timelapse recordings were captured continuously at 1 frame per second (fps) for 48 hours under constant temperature (18°C).

### Computational video processing and pose estimation

To track individual animal trajectories, raw time-lapse videos recorded at 8-hour increments were preprocessed. The video files were cropped into single-lane regions of interest corresponding to each animal. Coordinate boundaries were obtained using Fiji (ImageJ), and video cropping was executed using a custom Python script.

SLEAP (Social LEAP Estimates Animal Pose, (Pereira et al., 2022) was used for pose estimation. The model tracked three coordinates: the basal disk (foot), the center of the body column (body), and the hypostome (head). Additionally, the left and right boundaries of each microfluidic chamber were annotated as reference points for pixel-to-millimeter conversion. Following model predictions, mislabeled frames were manually corrected. All frames within the first 8 hours or until positive phototaxis were fully annotated to capture phototactic behavior, while one frame was annotated for every hour across the full 48-hour timeline to capture macro foot position. To reduce redundancies caused by the hydras’ slow movement and resting periods, coordinates were down sampled and extracted at 1 datapoint every 20 frames in Python for macro-kinematic behaviors.

### Behavioral analysis

Custom MATLAB code was used for data analysis and modeling. Foot coordinates were used to determine hydra location at a given time. The head orientation (θ) was calculated as the angle between the two vectors: (1) the vector connecting the body to the hypostome, and (2) the vector connecting the foot to the light source. Jump distance was calculated as the distance between the foot locations before and after a jump event. The number of jumps is the number of jump events observed.

To quantify postural persistence, we calculated the circular autocorrelation of head orientation across shifting time lags (k) using the mean cosine of angular differences: 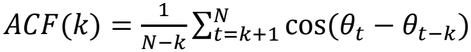 Postural half-life (τ) was defined as the lag time at which autocorrelation decayed below 0.5. For instances where autocorrelation did not drop below 0.5 within the maximum evaluated lag window of 133.3 minutes (n = 2), the half-life was conservatively assigned the maximum lag duration.

### Biased Correlated Random Walk Model

To simulate spatial trajectories, we adapted the Biased Correlated Random Walk (BCRW) model described by Kim et al. (2024). Because empirical head angle adjustments (Δθ) occur incrementally, sequential head orientation updates (θ) were calculated by combining the target direction (Ω_T_=180°) and the previous heading (θ_t_), weighted by a bias parameter *w* and persistence parameter (*1 - w*):

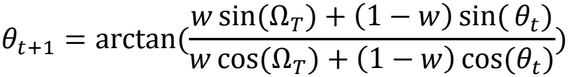

where δ_t_ represents stochastic angular noise sampled from a truncated location-scale distribution fitted to empirical data.

Bias parameter was calculated as:

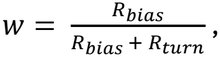

where,

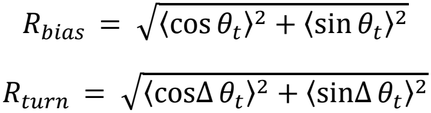

### Statistical analysis

The circular-analog of the Kruskal–Wallis test for directional data (Circstat MATLAB package; <u>Berens, 2009</u>) was used to evaluate the statistical significance of head orientations. The Mann–Whitney *U*-test (Wilcoxon rank sum test) was used to evaluate the statistical significance of jump distances, number of jumps, head autocorrelation, time spent in behavior, and transitions between behaviors. The Wilcoxon signed-rank test was used to evaluate the statistical significance of jump directions.

### Tracking Mask

Tracked coordinates for the head, body, and foot were filtered and cleaned independently. Pose estimates were classified as low confidence if confidence scores fell below 0.30 for the head and foot, or below 0.20 for the body (which exhibited lower baseline scores across postural configurations). Manual ground-truth annotations were held constant and excluded from filtering or interpolation.

Suspect frames (missing, low-confidence, or removed excursions) occurring in contiguous runs of 20 frames (0.67s) bound by valid coordinates were reconstructed using piecewise cubic Hermite interpolating polynomials (pchip in MATLAB) across x and y axes independently. This non-overshooting method prevented the creation of false movement artifacts. Gaps exceeding 20frames or contacting recording boundaries were blanked. The 20-frame interpolation threshold was empirically validated by reconstructing artificially hidden ground-truth coordinates, yielding a median spatial error of 0.24mm across 20-frame gaps. Across all animals, a median of 4.9% of frames were interpolated, 1.4% were removed as excursions, and 0.5% were blanked. High-frequency spatial noise (“chatter”) was flagged whenever a point’s coordinate deviated >1.9mm from its centered 61-frame rolling median. Chatter was maintained strictly as a diagnostic quality flag rather than an exclusion filter to prevent the accidental deletion of high-velocity biological events such as jumps.

### Behavioral State Classification

For each hydra, we compute eight features to capture their posture and movement. These features include speed of the foot, body, and hypostome; rate of change in body length; bending rate; body-axis rotation rate; rate of change for head angle; body length relative to its median. We compute the mean and standard deviations of these features on both short, medium, and long timescales to capture behavioral bouts that may vary from a few seconds to a few minutes long.

We used this feature set to train individual L1-regularized logistic regression models to predict hydra behavior (stillness, elongation, contraction, bend/sway). We hand-labeled behaviors from 30 no-light and start-dark hydra, and the L1 regularization ensured we penalized features that hurt the model or were redundant. This was important to prevent overtraining the model to a specific animal and allowing a trained model to be transferred zero-shot. The final model was chosen via 5-fold cross-validation. We tested accuracy using leave-one-hydra-out validation. To estimate the level of chance in each computed accuracy, we computed the shuffled null (**Figure 4–figure supplement 3**). Per-second predictions flicker, as features for each model may overlap, so we smoothed them into behavioral bouts, based on typical bout lengths seen in the hand labels. Light and dark animals were compared with two-sided Mann– Whitney U tests, with each animal as one data point.

## Data Availability

All raw and processed *Hydra* tracking trajectories, extracted kinematic features, and manual behavioral annotations are deposited on Zenodo under a CC BY 4.0 license (https://doi.org/10.5281/zenodo.24329561). Biased correlated random walk (BCRW) simulation code and the behavioral classifier code are available on GitHub (https://github.com/gbfranco/hydra-phototaxis) under an MIT license.

## Notes

https://github.com/gbfranco/hydra-phototaxis

https://doi.org/10.5281/zenodo.24329561

### Competing Interest Statement

The authors have declared no competing interest.

### Funding

- W.M. Keck Foundation Medical Research Grant Program

## References

Ayroles, J. F., Buchanan, S. M., O’Leary, C., Skutt-Kakaria, K., Grenier, J. K., Clark, A. G., Hartl, D. L., & De Bivort, B. L. (2015). Behavioral idiosyncrasy reveals genetic control of phenotypic variability. Proceedings of the National Academy of Sciences, 112(21), 6706–6711. 10.1073/pnas.1503830112

Badhiwala, K. N., Gonzales, D. L., Vercosa, D. G., Avants, B. W., & Robinson, J. T. (2018). Microfluidics for electrophysiology, imaging, and behavioral analysis of Hydra. Lab on a Chip, 18(17), 2523–2539. 10.1039/C8LC00475G

Badhiwala, K. N., Primack, A. S., Juliano, C. E., & Robinson, J. T. (2021). Multiple neuronal networks coordinate Hydra mechanosensory behavior. eLife, 10, e64108. 10.7554/eLife.64108

Bierbach, D., Laskowski, K. L., & Wolf, M. (2017). Behavioural individuality in clonal fish arises despite near-identical rearing conditions. Nature Communications, 8(1), 15361. 10.1038/ncomms15361

Bosch, T. C., Krylow, S. M., Bode, H. R., & Steele, R. E. (1988). Thermotolerance and synthesis of heat shock proteins: These responses are present in Hydra attenuata but absent in Hydra oligactis. Proceedings of the National Academy of Sciences, 85(21), 7927–7931. 10.1073/pnas.85.21.7927

Buchanan, S. M., Kain, J. S., & De Bivort, B. L. (2015). Neuronal control of locomotor handedness in Drosophila. Proceedings of the National Academy of Sciences, 112(21), 6700–6705. 10.1073/pnas.1500804112

Cazet, J. F., Siebert, S., Little, H. M., Bertemes, P., Primack, A. S., Ladurner, P., Achrainer, M., Fredriksen, M. T., Moreland, R. T., Singh, S., Zhang, S., Wolfsberg, T. G., Schnitzler, C. E., Baxevanis, A. D., Simakov, O., Hobmayer, B., & Juliano, C. E. (2023). A chromosome-scale epigenetic map of the Hydra genome reveals conserved regulators of cell state. Genome Research, 33(2), 283–298. 10.1101/gr.277040.122

Chapman, J. A., Kirkness, E. F., Simakov, O., Hampson, S. E., Mitros, T., Weinmaier, T., Rattei, T., Balasubramanian, P. G., Borman, J., Busam, D., Disbennett, K., Pfannkoch, C., Sumin, N., Sutton, G. G., Viswanathan, L. D., Walenz, B., Goodstein, D. M., Hellsten, U., Kawashima, T., … Steele, R. E. (2010). The dynamic genome of Hydra. Nature, 464(7288), 592–596. 10.1038/nature08830

Dupre, C., & Yuste, R. (2017). Non-overlapping Neural Networks in Hydra vulgaris. Current Biology, 27(8), 1085–1097. 10.1016/j.cub.2017.02.049

Giez, C., Noack, C., Sakib, E., Hofacker, L.-M., Repnik, U., Bramkamp, M., & Bosch, T. C. G. (2024). Satiety controls behavior in Hydra through an interplay of pre-enteric and central nervous system-like neuron populations. Cell Reports, 43(6), 114210. 10.1016/j.celrep.2024.114210

Han, S., Taralova, E., Dupre, C., & Yuste, R. (2018). Comprehensive machine learning analysis of Hydra behavior reveals a stable basal behavioral repertoire. eLife, 7, e32605. 10.7554/eLife.32605

Harel, Y., Nasser, R. A., & Stern, S. (2024). Mapping the developmental structure of stereotyped and individual-unique behavioral spaces in C. elegans. Cell Reports, 43(9), 114683. 10.1016/j.celrep.2024.114683

Honegger, K., & De Bivort, B. (2018). Stochasticity, individuality and behavior. Current Biology, 28(1), R8–R12. 10.1016/j.cub.2017.11.058

Kepner, Wm. A., & Hopkins, D. L. (1924). Reactions of Hydra to chloretone. Journal of Experimental Zoology, 38(4), 437–448. 10.1002/jez.1400380403

Kim, S., Badhiwala, K. N., Duret, G., & Robinson, J. T. (2024). Phototaxis is a satiety-dependent behavioral sequence in Hydra vulgaris. Journal of Experimental Biology, 227(18), jeb247503. 10.1242/jeb.247503

Lenhoff, H. M. (1961). Activation of the Feeding Reflex in Hydra littoralis. The Journal of General Physiology, 45(2), 331–344. 10.1085/jgp.45.2.331

Linneweber, G. A., Andriatsilavo, M., Dutta, S. B., Bengochea, M., Hellbruegge, L., Liu, G., Ejsmont, R. K., Straw, A. D., Wernet, M., Hiesinger, P. R., & Hassan, B. A. (2020). A neurodevelopmental origin of behavioral individuality in the Drosophila visual system. Science, 367(6482), 1112–1119. 10.1126/science.aaw7182

Macias-Muñoz, A., Murad, R., & Mortazavi, A. (2019). Molecular evolution and expression of opsin genes in Hydra vulgaris. BMC Genomics, 20(1), 992. 10.1186/s12864-019-6349-y

Mast, S. O. (1903). REACTIONS TO TEMPERATURE CHANGES IN SPIRILLUM, HYDRA, AND FRESH-WATER PLANARIANS. American Journal of Physiology-Legacy Content, 10(4), 165–190. 10.1152/ajplegacy.1903.10.4.165

Moiron, M., Laskowski, K. L., & Niemelä, P. T. (2020). Individual differences in behaviour explain variation in survival: A meta-analysis. Ecology Letters, 23(2), 399–408. 10.1111/ele.13438

Morris Little, H., Primack, A. S., Tsverov, J., Nunneley, M. P., Mühlbauer, S., Cox, B. D., Busse, C., Schneid, S., Louwagie, A., Cazet, J. F., David, C. N., Farrell, J. A., & Juliano, C. E. (2025). A molecular, spatial and regulatory atlas of the Hydra vulgaris nervous system. Development, 152(20), dev204983. 10.1242/dev.204983

Passano, L. M., & McCullough, C. B. (1962). THE LIGHT RESPONSE AND THE RHYTHMIC POTENTIALS OF HYDRA. Proceedings of the National Academy of Sciences, 48(8), 1376–1382. 10.1073/pnas.48.8.1376

Pereira, T. D., Tabris, N., Matsliah, A., Turner, D. M., Li, J., Ravindranath, S., Papadoyannis, E. S., Normand, E., Deutsch, D. S., Wang, Z. Y., McKenzie-Smith, G. C., Mitelut, C. C., Castro, M. D., D’Uva, J., Kislin, M., Sanes, D. H., Kocher, S. D., Wang, S. S.-H., Falkner, A. L., … Murthy, M. (2022). SLEAP: A deep learning system for multi-animal pose tracking. Nature Methods, 19(4), 486–495. 10.1038/s41592-022-01426-1

Plachetzki, D. C., Fong, C. R., & Oakley, T. H. (2012). Cnidocyte discharge is regulated by light and opsin-mediated phototransduction. BMC Biology, 10(1), 17. 10.1186/1741-7007-10-17

Rushforth, N. B., Burnett, A. L., & Maynard, R. (1963). Behavior in Hydra: Contraction Responses of Hydra pirardi to Mechanical and Light Stimuli. Science, 139(3556), 760–761. 10.1126/science.139.3556.760

Scappaticci, A. A., Kahn, F., & Kass-Simon, G. (2010). Nematocyst discharge in Hydra vulgaris: Differential responses of desmonemes and stenoteles to mechanical and chemical stimulation. Comparative Biochemistry and Physiology Part A: Molecular & Integrative Physiology, 157(2), 184–191. 10.1016/j.cbpa.2010.06.177

Schroeder, L. A., & Callaghan, W. M. (1981). Thermal tolerance and acclimation of two species of Hydra 1. Limnology and Oceanography, 26(4), 690–696. 10.4319/lo.1981.26.4.0690

Singer, R. H., Rushforth, N. B., & Burnett, A. L. (1963). The photodynamic action of light on Hydra. Journal of Experimental Zoology, 154(2), 169–173. 10.1002/jez.1401540204

Trembley, A. (1744). Mémoires pour servir à l’histoire d’un genre de polypes d’eau douce, à bras en forme de cornes. Jean & Herman Verbeek.

Tzouanas, C. N., Kim, S., Badhiwala, K. N., Avants, B. W., & Robinson, J. T. (2021). Hydra vulgaris shows stable responses to thermal stimulation despite large changes in the number of neurons. iScience, 24(6), 102490. 10.1016/j.isci.2021.102490

Wagner, G. (1905). On Some Movements and Reactions of Hydra. Journal of Cell Science, S2-48(192), 585–622. 10.1242/jcs.s2-48.192.585

Yamamoto, W., & Yuste, R. (2023). Peptide-driven control of somersaulting in Hydra vulgaris. Current Biology, 33(10), 1893–1905.e4. 10.1016/j.cub.2023.03.047

