## Supplemental Figures for "Diverging light-responsive behaviors in genetically similar strains of *Hydra vulgaris*"

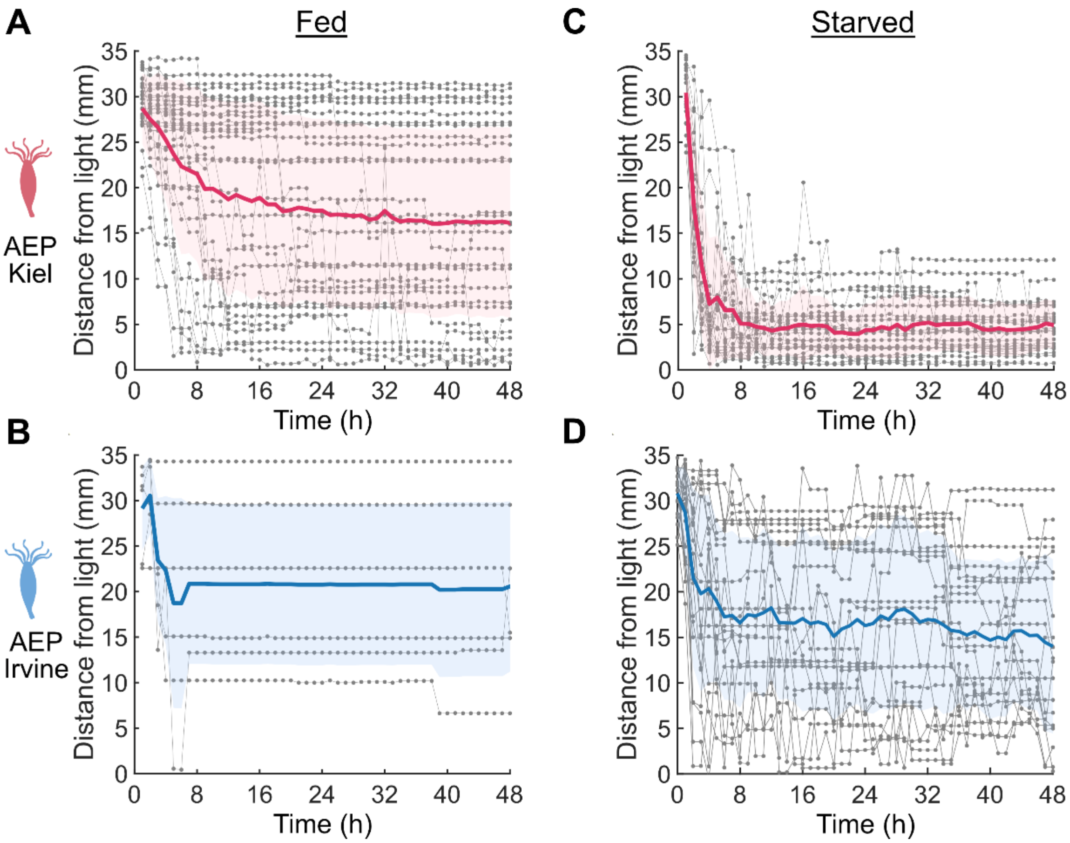


**Figure 1– figure supplemental 1. Foot position of *Hydra vulgaris lines* derived from AEP background over 48 h under light stimulus.** **(A–D)** Foot position relative to light (mm) sampled at every hour over 48 hours (t = 0, 1, ..., 48 h) under continuous directional light. **(A)** Fed AEP Kiel (n = 29) foot position plateaus near center of chamber after 8 h. **(B)** Starved AEP Kiel (n = 24) foot position moves towards light within 8 h and remains there. **(C)** Fed AEP Irvine (n = 6) foot position plateaus near center of chamber after 8 h. **(D)** Starved AEP Irvine (n = 21) foot position similarly plateaus near center of chamber after 8 h. Bold line and shaded band are group mean ± SD, and in gray are individual traces.

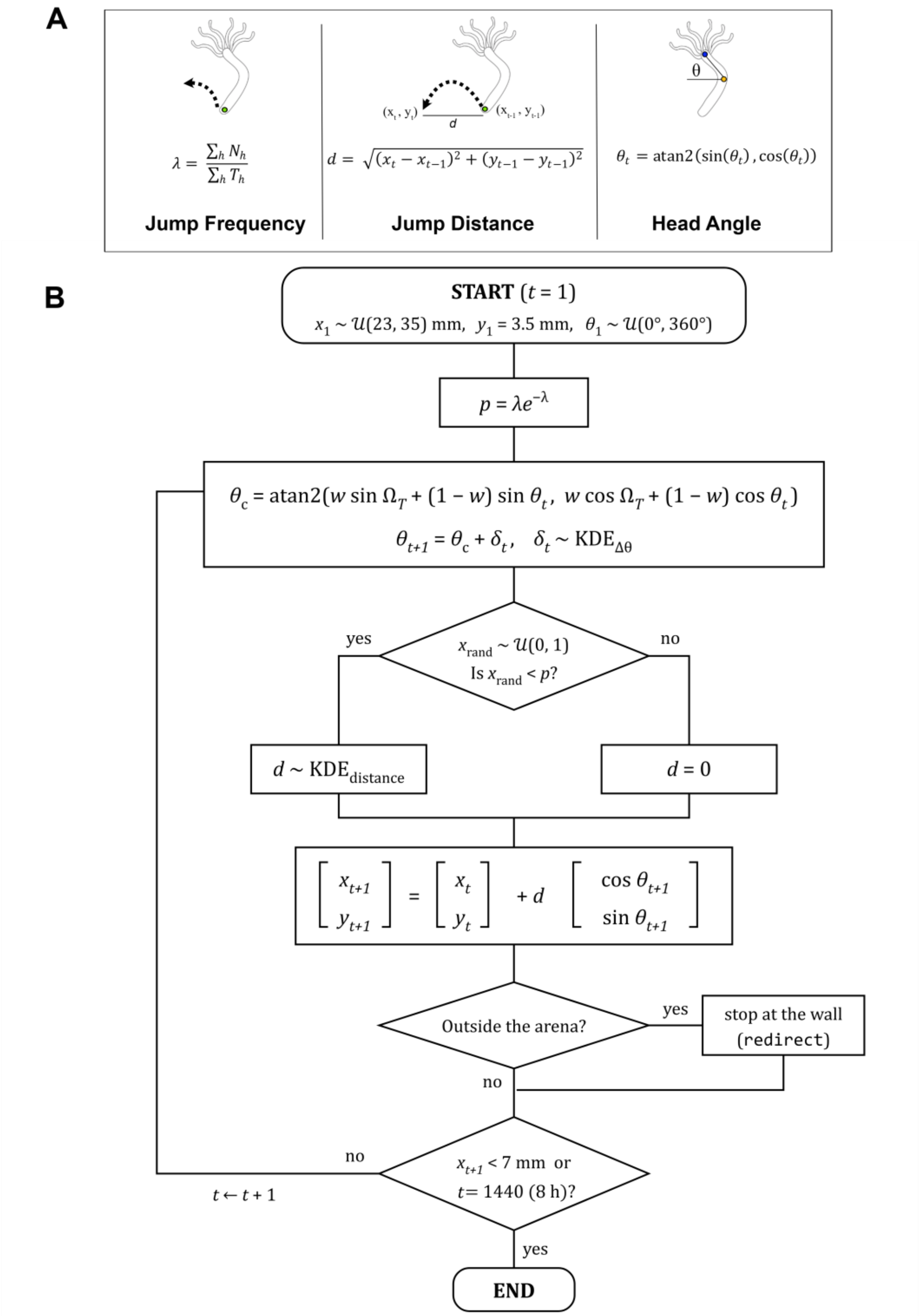


**Figure 3– figure supplement 1. Computational architecture of the Biased Correlated Random Walk (BCRW) model.**

**(A)** Schematic and mathematical formulations of empirical parameters extracted from tracking data to seed the BCRW model. *Left:* Jump frequency ($\lambda$), calculated as the mean rate of jump events across $H$ individual *Hydra* polyps, where $N_h$ is total jump count and $T_h$ is observation duration for *Hydra* $h$. *Middle:* Jump distance ($d$), defined as the Euclidean displacement between spatial coordinates at successive step intervals $(x_{t-1}, y_{t-1})$ and $(x_t, y_t)$. *Right:* Head angle ($\theta_t$), defined as the relative change in heading orientation $(\theta_t - \theta_{t-1})$ between consecutive frames. **(B)** Flowchart of the generative BCRW simulation algorithm. At step $t = 1$, agent spatial position is initialized in the far arena region ($x_1 \sim \mathcal{U}(23,35)$ mm, $y_1 = 3.5$ mm) with a uniformly distributed heading angle $\theta_1 \sim \mathcal{U}(0°, 360°)$. At each time step, stepwise jump probability $p = \lambda e^{-\lambda}$ and bias weight $w = \frac{R_{\text{bias}}}{R_{\text{bias}} + R_{\text{turn}}}$ (where $R = |\langle e^{i\theta} \rangle|$ represents mean resultant vector length) are evaluated. Updated heading direction $\theta_{t+1}$ is computed by vector-combining phototactic target direction ($\Omega_T$) and persistence heading ($\theta_t$) weighted by $w$, with stochastic turn noise $\delta_t$ sampled from the empirical turning angle kernel density estimate ($\mathrm{KDE}_{\Delta\theta}$). A uniform random draw $x_{\text{rand}} \sim$

$\mathcal{U}(0,1)$ determines jump execution: if $x_{\text{rand}} < p$, displacement $d$ is sampled from $\text{KDE}_{\text{distance}}$, otherwise $d = 0$. Position coordinates $[x_{t+1}, y_{t+1}]^T$ are updated via trigonometric projection, incorporating arena wall redirection rules when boundary contact occurs. Iterations ($t \leftarrow t + 1$) continue until reaching target threshold ($x_{t+1} < 7$ mm) or simulation time limit ($t = 8$ h).

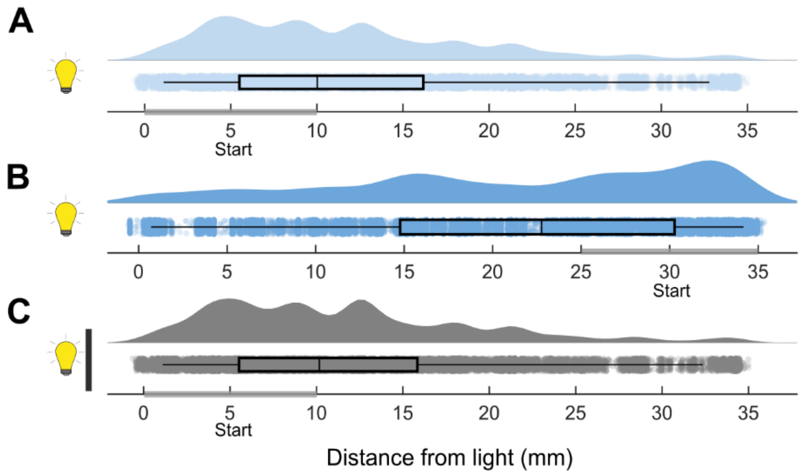


**Figure 4—figure supplement 1. Foot distribution of AEP Irvine hydras across starting positions relative to the light source. (A)** Animals were placed on the side nearest the light source (0–10 mm; n = 21). **(B)** Animals were placed furthest from the light source (25–35 mm; n = 21). **(C)** Animals were placed on the side nearest the light source with the light blocked (0-10 mm; n = 15). All box plots show medians, first, and third quartiles.

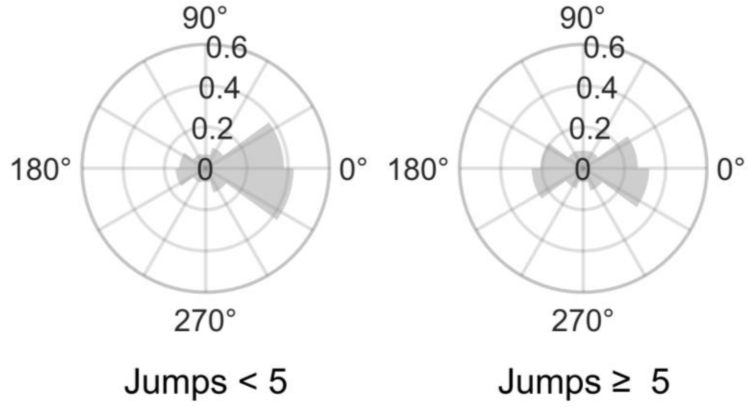


**Figure 4—figure supplement 2. Head angle of starved AEP Irvine *Hydra* in the absence of light split into groups based on jumps.** Left: Inactive animals (< 5 jumps) display head angles ($\theta$) oriented towards opposite side of initial placement position. Right: Active animals (≥ 5 jumps) exhibit uniform distribution of head angles.

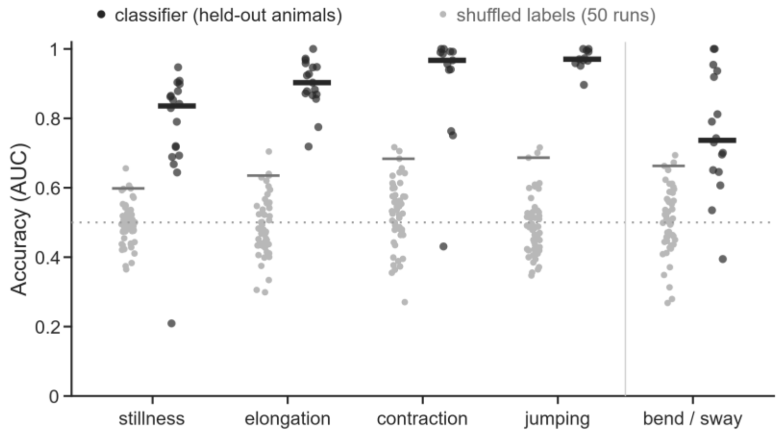


**Figure 4—figure supplement 3. Classifier accuracy and performance validation across behaviors.** Classifier accuracy (Area Under the ROC Curve, AUC) for predicting five hand-labeled *Hydra* behaviors (stillness, elongation, contraction, jumping, and bend / sway). Dark dots represent model performance on individual held-out animals evaluated using leave-one-hydra-out cross-validation. Light gray dots represent performance from shuffled-label null controls (50 runs). Horizontal dark bars indicate median AUC for held-out animals; light gray horizontal bars represent median AUC of the shuffled null baseline. The horizontal dotted line at $0.5$ denotes theoretical chance-level accuracy (AUC $= 0.5$).

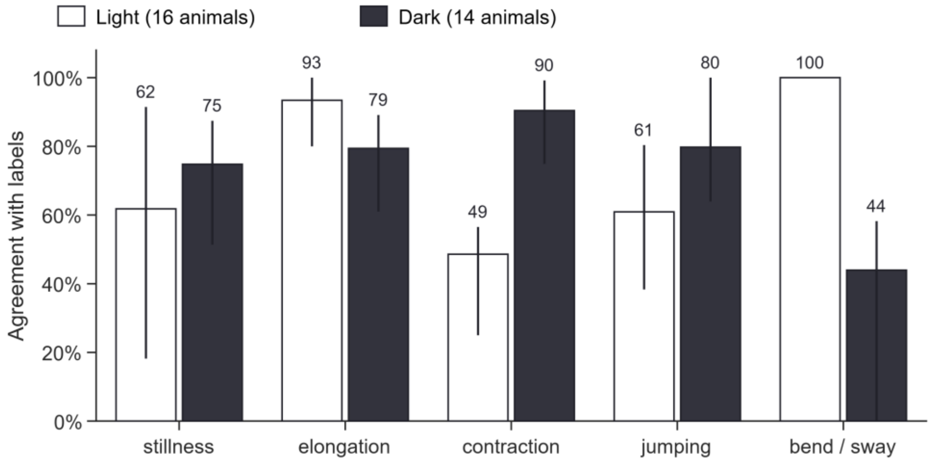


**Figure 4—figure supplement 4. Classifier agreement with manual behavioral annotations.** Frame-by-frame percentage agreement between automated classifier predictions and hand-annotated ground-truth labels across four behavioral categories (stillness, elongation, contraction, and jumping). Bars indicate total pooled agreement across all analyzed seconds; vertical error lines denote the interquartile range (25th–75th percentiles) across individual animals. Data are evaluated across three lighting regimes: Light (n = 16; unseen held-out test cohort), and dark (n = 14, white; training cohort). Numbers above each bar indicate the pooled agreement percentage. Discrepancies in the jumping category reflect differences between wider manually marked bout durations and point-event detection thresholds used by the classifier.

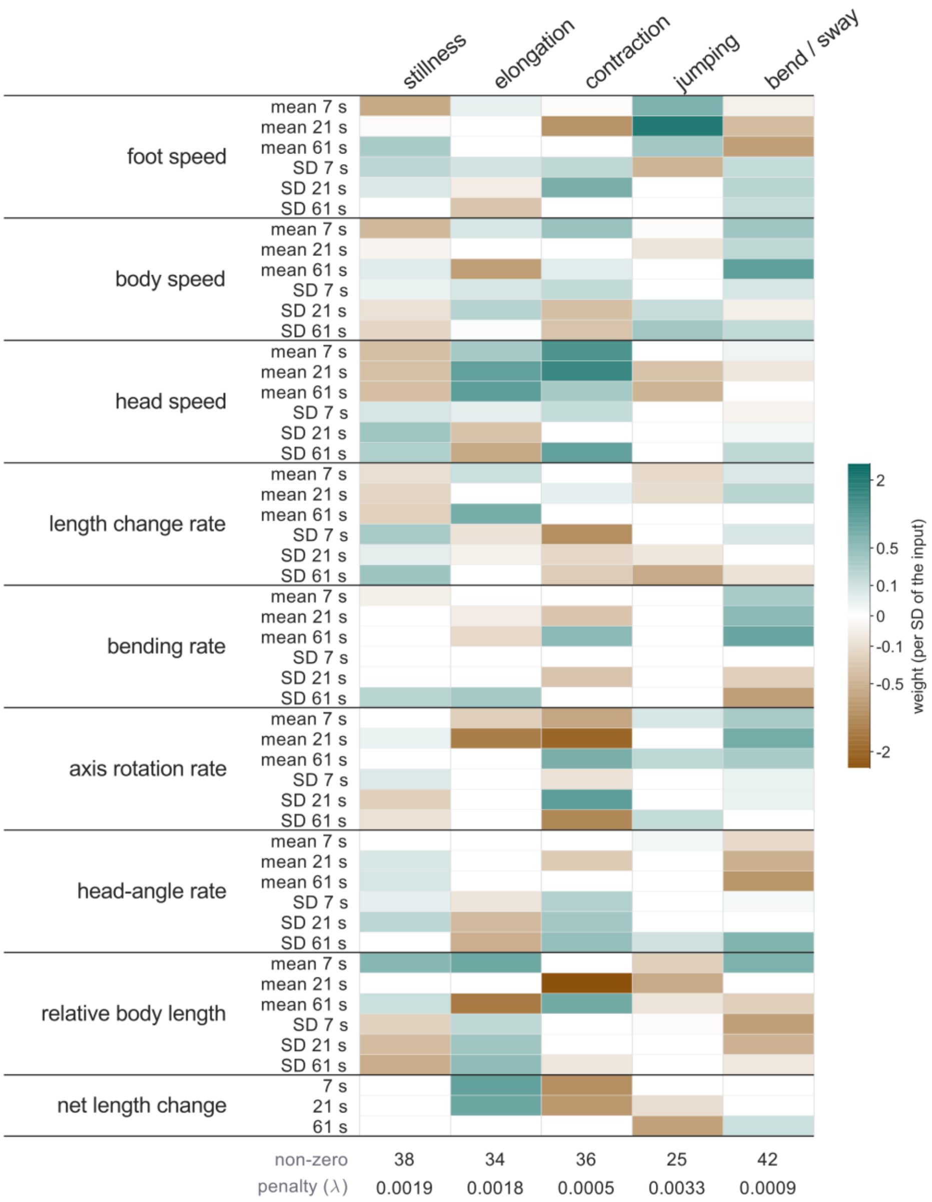


**Figure 4—figure supplement 5. Feature weights of L1-regularized logistic regression models across behavioral states.** Heatmap displaying normalized feature weights (coefficients) assigned by L1-regularized logistic regression models trained to classify five distinct *Hydra* behaviors (stillness, elongation, contraction, jumping, and bend / sway). Input kinematic features (rows) comprise rolling temporal means and standard deviations (SD) calculated across three sliding time windows (7 s, 21 s, and 61 s) for nine behavioral metrics (e.g., foot speed, head speed, axis rotation rate, relative body length). Color intensity represents weight per standard deviation of the input variable, ranging from positive predictors (teal) to negative predictors (brown); white cells indicate redundant or uninformative features assigned a weight of zero by L1 regularization. Bottom rows indicate the number of non-zero features retained (non-zero) and the optimal L1 regularization penalty (penalty (λ)) determined via 5-fold cross-validation for each binary classifier.

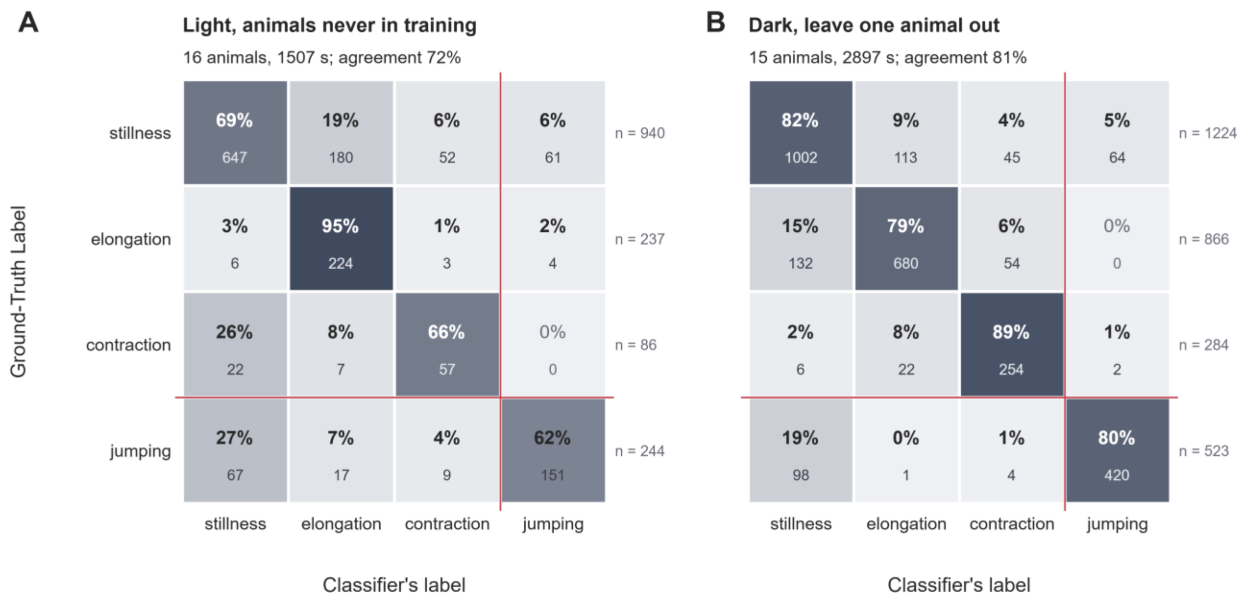


**Figure 4—figure supplement 6. Automated classifier accuracy and confusion matrix across experimental conditions. A)** Generalization test on held-out animals never exposed during training (Light condition; 16 animals, 1,507 s evaluated; 72% overall agreement). **B)** Leave-one-animal-out cross-validation (Dark condition; 15 animals, 2,897 s evaluated; 81% overall agreement). Numbers inside cells denote percentage of ground-truth seconds assigned to each predicted class (bold) and total duration in seconds (gray). Jumping (red) is hand-verified rather than predicted, so its row and column compare two hand passes; agreement over the three predicted behaviors is 77% (A) and 84% (B).
